# Social predation by a nudibranch mollusc

**DOI:** 10.1101/2024.07.01.600874

**Authors:** Kate Otter, Saida Gomidova, Paul S. Katz

**Affiliations:** Neuroscience and Behavior Graduate Program, University of Massachusetts Amherst, Amherst MA, USA; Department of Biology, University of Massachusetts Amherst, Amherst MA, USA; Neuroscience and Behavior Graduate Program and Department of Biology, University of Massachusetts Amherst, Amherst MA, USA

**Keywords:** foraging behavior, Berghia stephanieae, producing, scrounging, satiety-dependence, dangerous prey

## Abstract

Social predation is a common strategy used by predators to subdue and consume prey. Animals that use this strategy have many ways of finding each other, organizing behaviors and consuming prey. There is wide variation in the extent to which these behaviors are coordinated and the stability of individual roles. This study characterizes social predation by the nudibranch mollusc, *Berghia stephanieae*, which is a specialist predator that eats only the sea anemone, *Exaiptasia diaphana*. A combination of experimental and modeling approaches showed that *B. stephanieae* does predate upon *E. diaphana* in groups. The extent of social feeding was not altered by length of food deprivation, suggesting that animals are not shifting strategies based on internal state. It was unclear what cues the individual *Berghia* used to find each other; choice assays testing whether they followed slime trails, were attracted to injured anemones, or preferred conspecifics feeding did not reveal any cues. Individuals did not exhibit stable roles, such as leader or follower, rather the population exhibited fission-fusion dynamics with temporary roles during predation. Thus, the *Berghia* provides an example of a specialist predator of dangerous prey that loosely organizes social feeding, which persists across hunger states and uses temporary individual roles; however, the cues that it uses for aggregation are unknown.

**Significance Statement:** Social predation is a strategy to hunt dangerous prey and minimize injury. Many nudibranchs specialize as predators of cnidarians, which are dangerous to them. Although nudibranchs are typically characterized as solitary hunters, we provide evidence that social predation strategies may be used by a species that specializes on one species of sea anemone. The study showed that the individual sea slugs assumed temporary roles for establishing groups and that the group dynamics were unstable. However, the cues that the nudibranchs use to aggregate remain elusive.

## Introduction

Social feeding behaviors are seen in across taxa from one of the simplest multicellular animals *Trichoplax adherans* to more complex animals such as cephalopods and wolves (Burford and Robison, 2020; Fortunato and Aktipis, 2019; MacNulty et al., 2014). Social predation includes a suite of behaviors that encompasses the species’ degree of sociality and dependence on social foraging as well as its communication, specialization, and resource sharing (Lang and Farine, 2017). These different traits can be found in various combinations depending on the species. Complex social predation strategies, which include choreographed attack patterns, are used by animals that live socially (Berghänel et al., 2022) as well as by animals that are generally solitary (Lührs and Dammhahn, 2010; Twining and Mills, 2021). Some species are able to subdue or consume their prey in a way that is more efficient as a result of their collective actions, for example electric eels sometimes herd and surround their prey and then subdue large numbers with joint electrical strikes (Bastos et al., 2021) and lionfish which have increased success during hunting when multiple individuals attack the same group, but do not show coordination (Lönnstedt et al., 2014; Sarhan and Bshary, 2023). This framework also encompasses more simple forms of social predation characterized by aggregation, such as that of brown bears who live solitarily, but aggregate at food sources (Deacy et al., 2016), and predatory nematode-hunting mites that aggregate around injured prey (Aguilar-Marcelino et al., 2014). Aggregation as a strategy differs from other strategies in that the individuals are feeding together but their behaviors are largely independent of one another. In this paper, we sought to characterize the social predation of a nudibranch that feeds on sea anemones and the cues that the animals use to aggregate.

This diversity of social predation strategies across taxa is influenced by the range of costs and benefits incurred during social predation and feeding. Hunting and foraging in groups can make it easier to find food because the load of searching and identifying a food source is split among more individuals (Snijders et al., 2021). However once located, the food must be shared across more individuals (Sutton et al., 2015). If animals experience predation risk, a benefit of social feeding is increased vigilance of the group with reduced energy spent on individual vigilance allowing more time to feed (Barta et al., 2004; Kelley et al., 2011). On the other hand, groups can make it easier for predators to spot and capture individuals (Balaban-Feld et al., 2019; Sutton et al., 2015). A key benefit of social predation is the capacity to subdue and kill larger, more dangerous prey (Brown and Richardson, 1988; MacNulty et al., 2014; Mukherjee and Heithaus, 2013). Hunting in groups can reduce the risk of injury for each individual and minimize the amount of handling time, which is when injury typically occurs.

The nudibranch, *Berghia stephanieae* is a monophagous specialist predator that feeds on a single species of sea anemone, *Exaiptasia diaphana* (Carroll and Kempf, 1990; Goodheart et al., 2022; Monteiro et al., 2020). Like other sea anemones, *E. diaphana* is dangerous due to its nematocysts and acontia, specialized structures for deterring predators; it can even kill and consume potential predators (Hayes and Schultz, 2022; Lam et al., 2017; Mehrotra et al., 2019). Thus, it is possible that *B. stephanieae* socially feeds as a strategy to minimize risk of severe injury. Here we sought first to establish that aggregation at anemones is not by random chance and the behavioral mechanisms that *Berghia* uses to decide when and where to feed. It is possible that *Berghia* follows the slime trails of conspecifics to anemones to aggregate. Nudibranchs, like other gastropods, rely on deposition of mucus that they glide on using cilia on their muscular foot. In terrestrial and aquatic gastropods, trail following is a mechanism that many species use to find mates (Ng et al., 2013), hunt other gastropods (Leonard and Lukowiak, 1984; Patel et al., 2014) and otherwise aggregate (Bretz and Dimock, 1983; Davies and Beckwith, 1999). Similarly, the marine gastropod, *Aplysia californica*, aggregates at conspecific odors (Audesirk, 1975).

Another potential mechanism is related to the concept of social influence (Webster and Fiorito, 2001; Whiten and Ham, 1992). Social influence is when the actions of conspecifics drive behavioral changes and/or shifts in motivational states of an individual. For example, some crabs locate food by observing other crabs eating; the presence of crabs eating acts to stimulate eating (Kurta, 1982). Similarly, meat traps for *Vespula germanica* wasps are facilitated by the presence of conspecifics at the trap (D’adamo et al., 2003).

Aggregation could also be caused by chemical cues emanating from injured prey. Kairomones from prey have been shown to attract predators in species such as nematode-hunting mites (Aguilar-Marcelino et al., 2014; Bilgrami, 1994), frogs (Schoeppner and Relyea, 2005; South et al., 2020). Similarly, herbivorous species such as silkworms are attracted to leaves with conspecific damage (Mooney et al., 2009). Alarm pheromones have been identified in some species of anemones (Howe and Harris, 1978). Thus, it is possible that *B. stephanieae* could detect and respond to such disturbance cues, alarm cues, or an increase in anemone volatiles released by the anemone during consumption by other slugs. In this study, we experimentally establish that *Berghia* feeds socially even with the opportunity to feed individually. We tested several different mechanisms that could be used by individuals to locate conspecifics during predation. We also tested whether *Berghia* changes its group feeding based on hunger state and whether individuals show consistent preferences to feed in groups.

## Methods

### Animal Care

A colony of *Berghia stephanieae* was maintained from individuals purchased from Salty Underground (Crestwood, MO, USA) and Reeftown (Boynton Beach, FL, USA). Prior to use in this study, *Berghia* were communally housed in groups of 5-15 individuals in 1-gallon acrylic aquariums filled with artificial seawater (ASW; Instant Ocean, Blacksburg, VA, USA), made with a specific gravity of 1.020 --1.022 and pH of 8.0 - 8.5 with a 12:12 light dark cycle at 22-26°C.

*Exaiptasia diaphana* (Carolina Biological Supply Co., Burlington, NC, USA) were housed in glass aquariums containing ASW. Unless otherwise noted, the *B. stephanieae* were fed twice a week by placing two *E. diaphana* individuals in their home tank.

### Group Feeding Assay

For the group feeding assay, eight *E. diaphana* individuals were placed in a circle around the edge of a large clear square acrylic box (25 × 25 × 25 cm). The arena was situated on a white LED lightboard with opaque black electrical tape around the outside edges to exclude visual stimuli. The animals were recorded from above using a Pro Stream Webcam 1080P HD at 1 FPS using Video Velocity software (Virginia City, Nevada, USA). The anemones were allowed to acclimate for 5 minutes and then eight *B. stephanieae* were added to the center of the circle. After 20 minutes the sizes of the groups and number of slugs that were not feeding were recorded and the slugs were returned to their home tanks. The slugs used in these experiments were food-deprived for either 7-days or 3-days depending on the experiment. Group sizes were counted by observers blind to the food-deprivation length.

### 2-Alternative Choice Assays

There were several 2-alternative choice assays (Table 1). Two anemones were placed into a small square acrylic arena (7.62cm X 7.62cm X 2.54cm) whose sides were covered with opaque white window film. The arena was placed on a white LED light board. The anemones were allowed to acclimate in the arena for 5 minutes before a *B. stephanieae* was introduced. The *B. stephanieae* were acclimated in an identical arena on the lightboard in ASW or anemone-treated water (ATW) depending on the experiment. The ATW was ASW that had previously contained one anemone per 25 mL for at least 24 hours prior to filtration through a 0.22 µm PES filter (Milipore Sigma, Burlington, MA). Results were compared to an intact anemone that had not interacted with a slug before the trial.

**Table 1.**
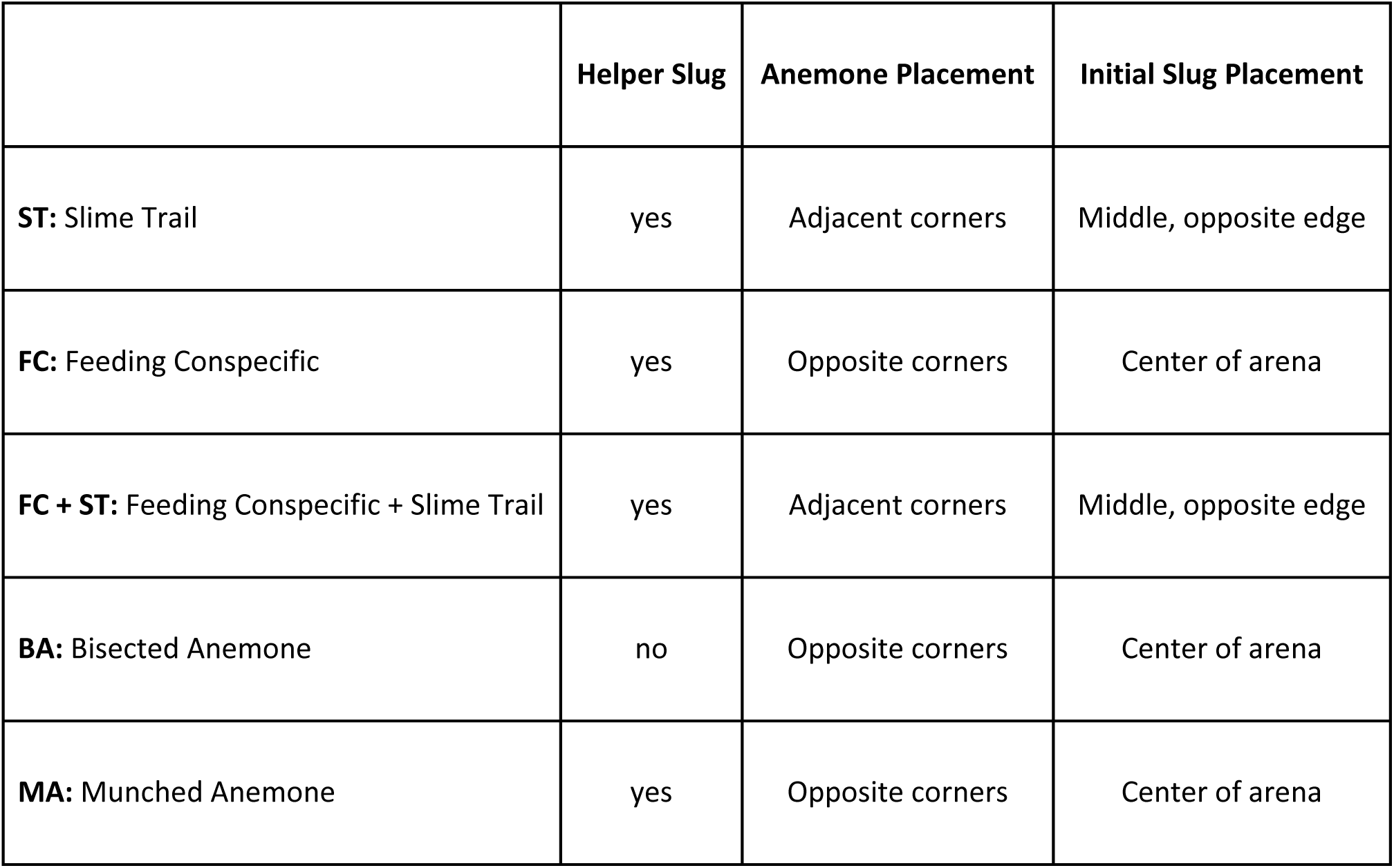
Descriptions of the 2-alternative choice assays.

Each trial was recorded from above using the same equipment and frame rate as the group feeding assay. The trial ended when the focal slug first made contact with one of the anemones. The data were not recorded blind, because the trials used a focal animal and the manipulation was visible to the observer. If the focal slug did not make contact with either anemone within 30 minutes, the trial was ended and omitted from subsequent analysis. For trials that involved a feeding conspecific, the trial was omitted if the helper slug stopped feeding before the focal slug selected an anemone. At the end of each trial the slugs were returned to their home tanks and their choice was recorded. All *Berghia* were food-deprived for 3 or 7 days depending on the experiment.

### Statistical Analysis

To statistically compare the group sizes observed with the null hypothesis that each slug chose independently of each other, we constructed a model with m slugs each selecting one of n anemones with equal probability (Eq. 1). Using this model, we simulated a trial and calculated the mean and maximum group sizes. This was repeated for the same number of trials in each dataset and then the mean of the mean and maximum group sizes were calculated for each simulated dataset to create the null distribution. The experimental means of the mean and maximum group sizes were then compared to the null distribution and the probability of the null model producing the same result or larger than the experimental data for a p-value was calculated. 100,000 datasets were simulated for each statistical test.

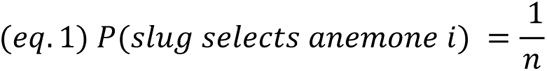

Additionally, we used the social dining model (SDM; often referred to as the “Chinese Restaurant Process”; (Antoniak, 1974; Pitman, 2002) to estimate a concentration parameter representing the propensity of individuals to select an anemone with feeding conspecifics. The social dining model is a discrete process that simulates a set of individuals, m, each sequentially selecting a dining location, m (Eq. 2). There is also a concentration parameter, α, that dictates how likely individuals are to select a dining location that is already occupied (Eq. 2). This model assumes that the number of anemones, n is greater or equal to the number of slugs, m. The model also assumes that the order in which the slugs choose does not affect the final probability distribution. We estimated the concentration parameter using a bisection method to iteratively determine the parameter that fits the experimental data. We used this parameterized model to calculate a p-value similarly to above.

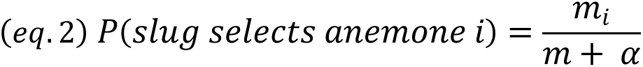

In addition to the models described above, the group feeding assay for 3- and 7-day food-deprived animals were compared using a t-test. For the 2-alternative choice assays, the proportion of individuals that selected the manipulated anemone was compared to random chance (50%) using a binomial proportion test.

The individual repeated measurements assay was analyzed using a Fisher’s Exact Test. We also tested if the distribution of social feeding scores (number of trials social option was selected) was bimodal using the Silverman (1981) critical bandwidth test as implemented by the *multimode* package (v1.5; Ameijeiras-Alonso et. al., 2021). To assess the individual repeatability of 2-alternative choice test outcomes, we estimated individual repeatability using the *rptR* package (v0.9.22; Stoffel et. al., 2017) with their choice in the predator-prey ratio assay as a predictor and individual identity as a random intercept.

All modeling, visualization and statistical analysis was performed in R version 4.2.3 (R Core Team 2023). Data manipulation used the *dplyr* (v1.1.4; Wickham et. al., 2023a) and *tidyr* (v1.3.0; Wickham et. al., 2023b) packages. For visualization we used the following packages: *ggplot2* (v3.4.4; Wickham, 2016), *ggpubr* (v0.6.0; Kassambara, 2023a), *rstatix* (v0.7.2; Kassambara, 2023b) and *cowplot* (v1.1.1; Wilke, 2020). All code to reproduce this analysis and the figures in this paper is available on Github.

## Results

### *Berghia* fed in groups more than expected by random chance

This study was inspired by observations of large groups of slugs forming during feeding in the laboratory (Fig. 1a). We quantified the distribution of slugs 20 minutes following a routine feeding and found that when fed with two anemones per tank, the slugs did not evenly distribute between the two anemones (Fig. 1b, c). A one-sample Wilcoxon signed-rank test indicated that the mean proportion of slugs feeding on one of the anemones, 0.85, was significantly different from 0.5 (Z = 231, p = 0.000052, effect size= 0.887).

**Fig. 1.**
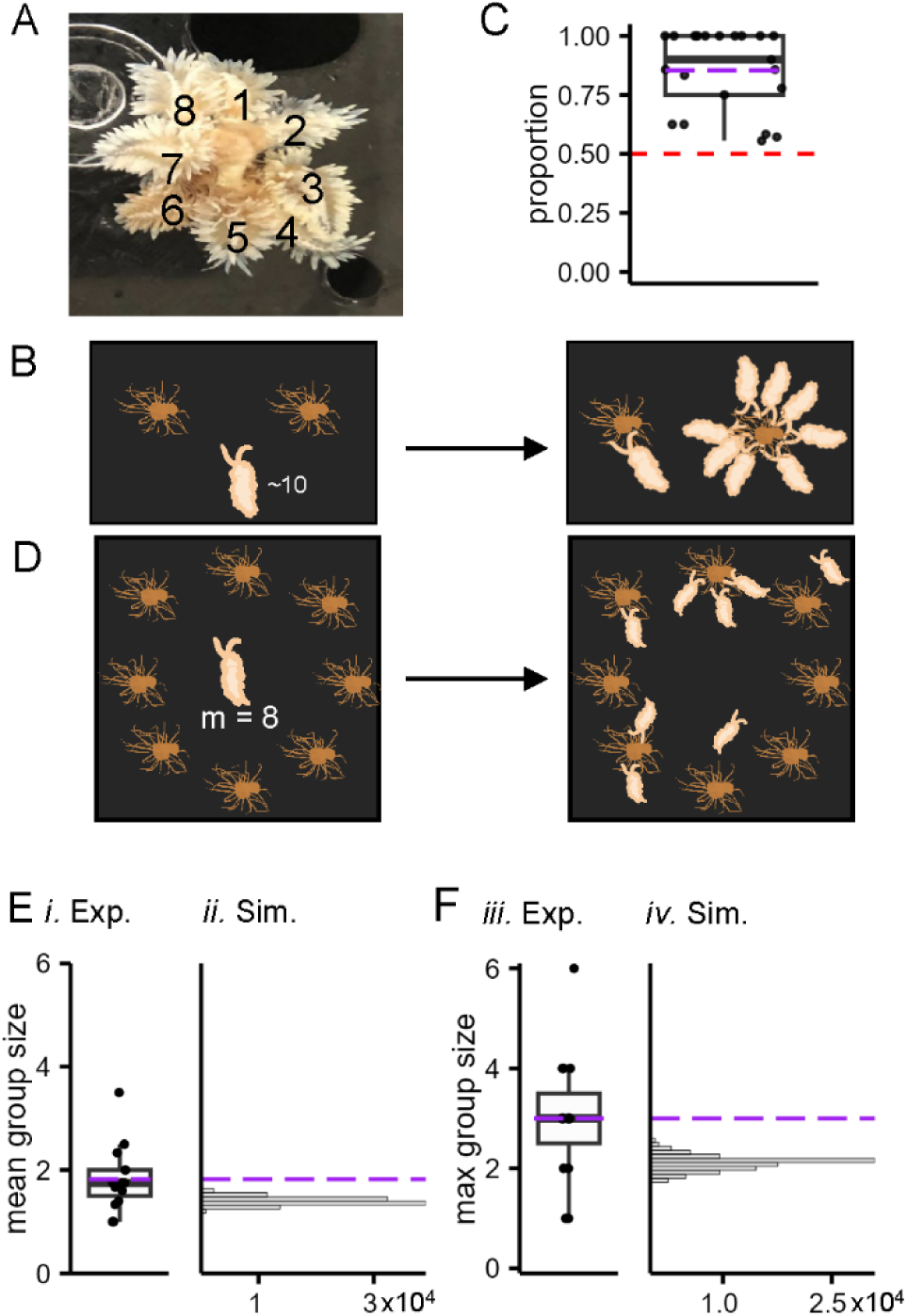
*Berghia stephanieae* form groups larger than if they each selected an anemone independently of each other. **A** Eight *B. stephanieae feeding on a single E. diaphana anemone* in an aquarium. The slugs are numbered for clarity. **B** A schematic showing the experiment used to quantify the grouping during routine feeding in their home tanks. Two anemones were placed into each home tank, which contained about ten slugs. After 20 minutes, the proportion of slugs feeding in the larger group was counted. **C** A boxplot of the proportion of slugs feeding on one of the two provided anemones. The slugs did not distribute evenly between the two anemones and tended to form large groups around one of them (Z = 231, p = 0.0000258). Purple line represents the mean. Red dashed line represents even distribution between the anemones. **D** A schematic of the group feeding assay (GF). **E** A boxplot representing the mean group sizes observed in each trial (left) and a histogram of the mean group sizes for each simulated dataset with the same number of trials as the experimental data of the null hypothesis where each slug selects an anemone independently of each other (right). The purple dashed line represents the mean group size of the experimental dataset. The observed mean does not occur within the distribution of the simulated data. **F** The same plots as **E**, using the maximum group size observed. Similarly, the observed mean of the max group sizes in the dataset does not occur in the simulated data. The simulated data sets have units of 10,000 datasets.

To test whether *B. stephanieae* feed on *E. diaphana* in groups even if they have the option to feed alone, we performed a group feeding assay. When given the opportunity to feed individually with a 1:1 ratio of *B. stephanieae* to *E. diaphana* (Fig. 1d), *Berghia* fed in groups larger than expected if each individual *B. stephanieae* was selecting an anemone independently of one another. Across 28 trials, the mean of the average group sizes observed in each trial was 1.82 (Fig. 1e*i*; median = 1.75, SD = ± 0.62) and the mean of the maximum group sizes observed in each trial was 3 (Fig. 1e*iii*; median = 3, SD = ± 1.25).

To distinguish an active choice to aggregate around prey from random grouping, we simulated a scenario where each slug selected an anemone with equal probability (eq. 1), which is representative of conditions where each individual slug was selecting prey independently of one another. 100,000 datasets with 28 trials each were simulated. There was no overlap between the experimental dataset mean and the simulation distribution; the experimental mean average group size was significantly more than expected by the simulated data (p = 0.00002; Fig. 1e*ii*) and the mean max group size of the experimental data was similarly larger than the simulated data (Fig. 1e*iv*; p = 0). Thus, the slugs are not choosing the anemones independently of each other.

### *Berghia* did not use the presence of feeding conspecifics or slime trails to select anemones to feed on

A series of 2-alternative choice assays were performed to examine potential cues that *B. stephanieae* could be using to aggregate (Table 1). One such cue is that the animals could be following the slime trail (ST) left by a conspecific animal. In the testing arena, a helper slug was placed in the middle of the arena and allowed to navigate the arena until it contacted one of two intact, size-matched anemones. Once the helper slug protruded its proboscis, it was removed from the arena before it could bite the anemone (Fig. 2A*i*). The target slug was then placed in the arena to determine which anemone it would choose, the one with the slime trail leading to it or the other. The slugs did not choose to feed on an anemone with a slime trail laid by a conspecific leading to it more than chance (p = 0.43, 23 out of 40). The slugs were also tested following an acclimation in ATW, to test whether prey scent would cause them to feed in groups due to heightened arousal. The target slug did not choose to feed on anemone with the slime trail leading it regardless of whether it was acclimated in ASW or in anemone-treated water (ATW) (Fig 2b; p = 0.61, 6 out of 15).

**Fig. 2.**
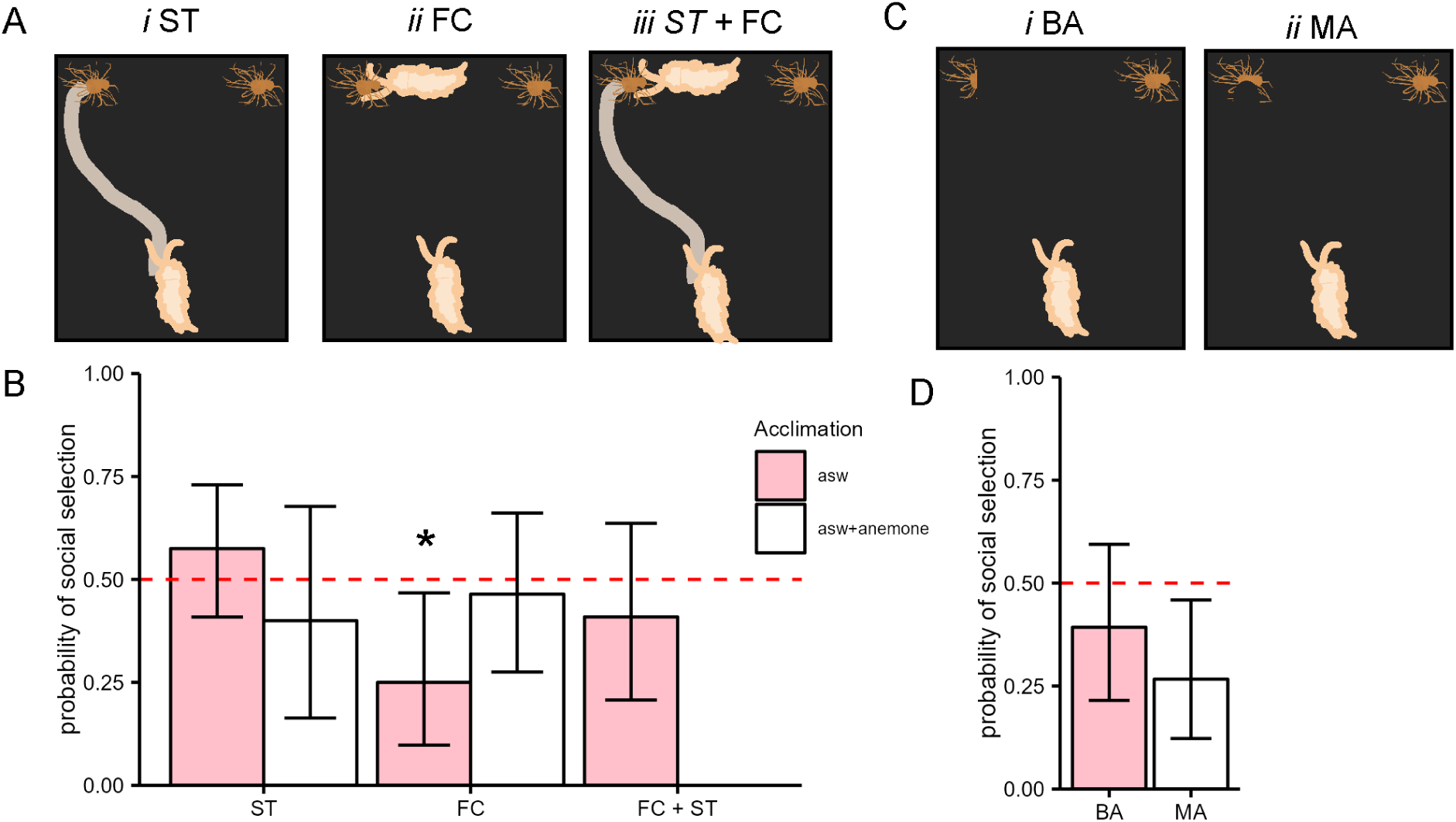
Behavior in 2-alternative choice tasks. **A** Schematics of the choice between an intact anemone and an anemone with a slime trail (ST), a feeding-conspecific (FC) or both (FC + ST). **B** Bar plots showing the proportion of animals that selected the manipulated anemone for each of the choices depicted in A. The red dashed line indicates random choice. Error bars represent 95% credible intervals of the binomial test. Pink bars represent slugs that were acclimated in filtered ASW and white bars represent slugs acclimated in anemone-treated water (ATW). All choices were not significantly different from random chance, except FC when acclimated in ASW which was selected lower than chance, meaning the slugs preferred an anemone without a feeding conspecific (8/30, p = 0.016). **C** A schematic of the choice between an intact anemone and an anemone that had been cut in half (BA) and an anemone that had been previously fed on by a conspecific (MA). **D** Bar plots showing the same as in **B**. None were significantly different from random chance.

To test whether slugs were simply attracted to a feeding conspecific (FC), a two choice test was constructed; a helper slug was placed in the center of a separate arena and allowed to begin feeding on one of two intact, size-matched anemones. Once the helper slug had chosen, both anemones were transferred to the testing arena with the helper slug still feeding on the anemone it had chosen. This avoided having a slime trail lead to the anemone. When given the choice to feed on an anemone with a feeding conspecific, the target slug actually preferred the intact anemone (Fig. 2b; p = 0.023, 6 out of 24). However, this preference went away when the target slug was acclimated in ATW (Fig. 2b; p = 0.85, 13 out of 28).

To determine whether the slugs needed the combination of a slime trail plus a feeding conspecific (ST+FC), a helper slug was placed in the middle of the testing arena and allowed to navigate the arena until it began feeding on one of two intact, size-matched anemones. The target slug was then placed in the arena. There was no preference shown for either anemone despite the presence of both a slime trail and feeding conspecific (Fig 2b; p = 0.52, 9 out of 22).

### *Berghia* did not prefer anemones that have been injured

The potential influence of kairomones from injured *E. diaphana* was tested with a 2-alternative choice assay. Two size-matched anemones were selected and one was cut in half with a razor blade (bisected anemone, BA). After 5 minutes, a target slug was added to the arena. *Berghia* showed no preference when given a choice between a bisected anemone and an intact anemone, (Fig 2d; p = 0.35, 11 out of 28).

To test whether slugs preferred an anemone that had been injured by a conspecific (munched anemone, MA), a helper slug was placed in the center of an arena and allowed to begin feeding on one of two intact, size-matched anemones. After 5 minutes of feeding the anemones were removed and placed in the testing arena. The target slug was then placed in the test arena. Surprisingly, slugs showed a preference for intact anemones over anemones that had been previously fed on by a conspecific (8/30, p =0.016; Fig. 2d).

Although the slugs did not show a preference to various social cues, they might have contacted the manipulated anemone more quickly, which could lead to aggregation. A three-way ANOVA was used to compare the effect of the slugs’ choices, the acclimation method, and the anemone manipulation on the time it took them to make a choice. The latency to choose was log-transformed to normalize. Slugs that selected the social option did not do in less time than animals that selected the control anemones for any of the treatments (Supplementary Figure S1, Supplementary Table S1). There was no effect of choice (F(1) = 1.113, p = 0.293), nor a statistically significant interaction effect (F(3,3) = 0.617, p = 0.605). Although was an effect of the assay type on the latency to select an anemone (F(3) = 8.851, p = 1.89e-05), this was not a main effect of interest (Supplementary Table S2). Slugs that were acclimated in ATW were faster to choose an anemone (F(1) = 18.578, p = 2.88e-05), likely due to heightened arousal from the scent of their prey prior to entering the arena, which differs from previous findings that food-deprived slugs in an empty arena move slower when presented with a food odor (Quinlan and Katz, 2023). This effect interacted significantly with their choice (F(1,1) = 5.864, p = 0.0166), such that their latency was impacted the most when the slugs selected control anemone and ATW acclimation caused them to choose faster. There was no interaction effect between acclimation and manipulation (F(1,3) = 0.734, p = 0.3931), nor was there a three-way interaction between the terms (F(1,1,3) = 1.658, p = 0.1998).

To test whether slugs preferentially selected the larger anemones, a nested ANOVA was used to compare the mean difference in the chosen anemone diameter from the anemone that was not chosen to 0 and test for an effect of anemone manipulation. The mean difference in anemone diameter between the anemone choices was not significantly different from 0 for any of the manipulations (Supplementary Figure S2; F(6) = 0.594, p = 0.735).

### Social predation was not facilitated by intermediate levels of food-deprivation

Animals might be changing their social feeding strategies because of a trade-off between food-acquisition and injury. To test the prediction that social predation is more prevalent in animals that are intermediately hungry, we compared 3-day and 7-day food-deprived animals in a group-feeding assay. Across 13 trials, the mean of the average group sizes observed per trial was 1.85 for the 3-day group and 1.82 for the 7-day food deprived group (Fig. 3a*i*; 3-day median = 1.67, SD = 0.75, 7-day median = 1.75, SD = 0.62). The mean maximum group size was 3.00 for both groups (Fig. 3b*i*; 3-day median = 3.00, SD = 1.25; 7-day median = 3.00, SD = 1.29). The mean group sizes for the 3-days food deprived animals were not significantly different from those of the 7-days food deprived animals (Fig. 3a; t = 0.11074, df = 23.553, p = 0.9128). The maximum group sizes were also not significantly different (Fig. 3b; t = 0, df = 25.201, p = 1).

**Fig. 3.**
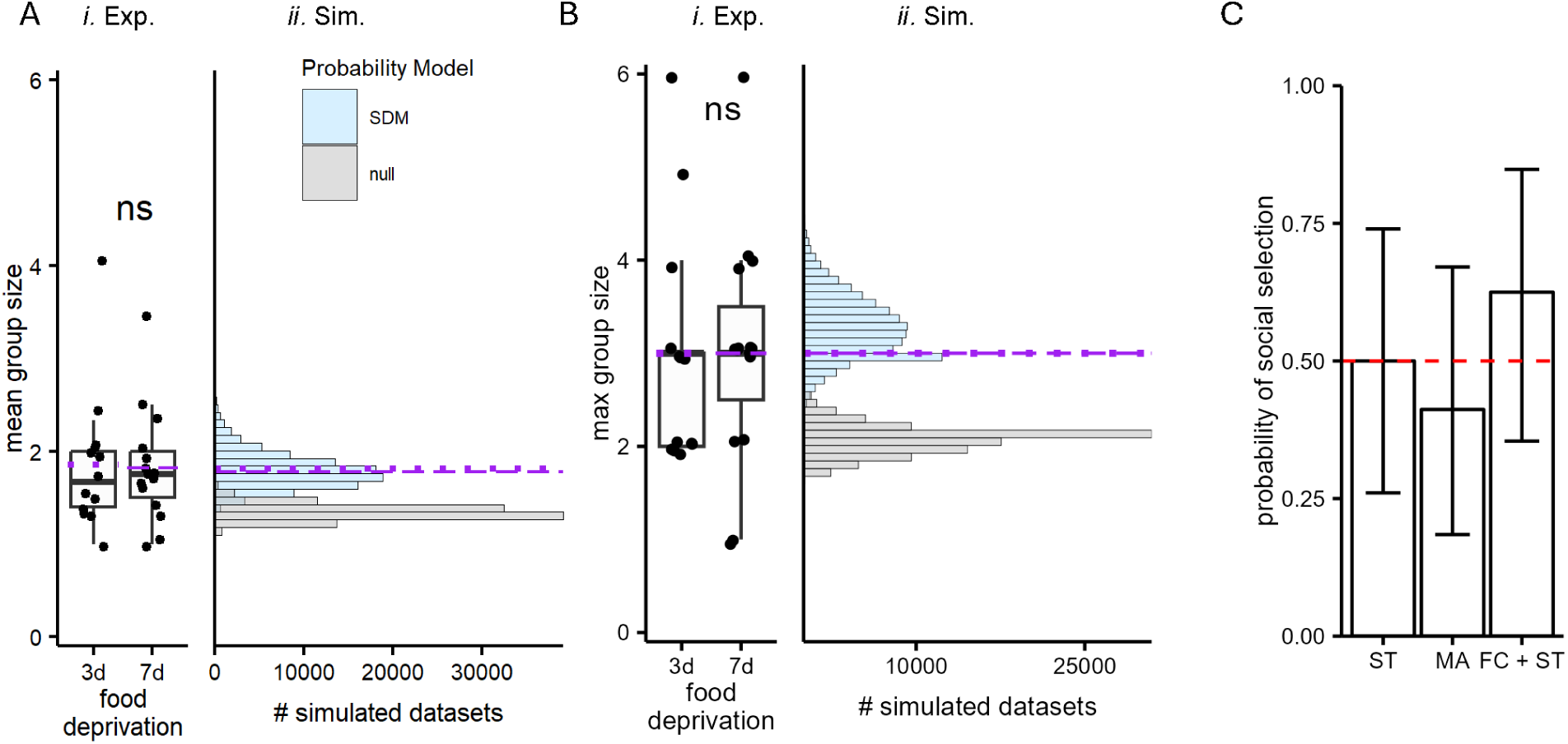
There is no difference in group size between intermediately food-deprived animals and 7-day food-deprived animals. **A** Boxplot of the mean group size for trials that were 3-days food-deprived and 7-days food-deprived (left). Histogram of the dataset mean of the mean group sizes observed in 10,000 simulated datasets (right). The light blue bars represent the parameterized social dining model (SDM), and the grey bars represent the null model. The dotted purple line is the experimental dataset mean for the 3-day food-deprived animals and the dashed purple line is the experimental mean for the 7-days food-deprived animals. There is no difference between the experimental means and they fall within the SDM simulated dataset means and do not overlap with the null simulated dataset means. **B** Similar plots as A for the dataset mean of the maximum group sizes. **C** The probability of selecting the manipulated anemone in 2-alternative choice assays comparing feeding conspecifics and slime trails (FC + ST), anemones previously fed on by a conspecific (MA) and anemones with slime trails (ST). None were significantly different from random chance (0.5).

Similarly to the 7-day food deprived data, we compared the 3-day food deprived experimental data to the distribution of simulated dataset means of the mean and max group size when each slug chose an anemone independently of one another with equal probability (eq. 1). These simulated datasets had 13 trials each, like the experimental dataset (Fig. 3a*ii*,b*ii*). The observed mean average group size and the mean maximum group size were significantly larger than the simulations (mean p = 0.00037, max p = 0.00084). Thus, 3-day food-deprived animals are also not choosing anemones independently of each other.

We also parameterized the social dining model (SDM) using the 3-days food-deprived dataset. The concentration parameter, α, was estimated to be 4.063. The distributions of the simulated dataset means for the mean and maximum group sizes included the experimental mean average group size (Fig. 3a*ii*; p = 1) and mean maximum group size (Fig. 3b*ii*; p = 1). Additionally, the 7-day food-deprived experimental mean of the mean and maximum group size was also within the simulated distribution parameterized with the 3-days food deprived dataset. This indicates that the grouping as indicated by the α parameter is similar for both levels of food-deprivation.

We also tested 3-days food-deprived animals in some of the 2-alternative choice assays. Like the 7-day food-deprived animals, 3-day food-deprived slugs showed no preference for any of the cues (Fig 3c). There were no differences between anemones with a slime trail (p = 1.00, 9 out of 18), anemones that were previously fed on by a conspecific (p = 0.63, 7 out of 17); or anemones with a feeding conspecific and a slime trail (p = 0.45, 10 out of 16).

### *Berghia* did not show consistent individual preferences to feed in groups

It is possible that the reason that no cue or hunger state was found to account for aggregation in social feeding could be that individual slugs have consistent preferences to feed socially or not. This individual preference might have been lost in the group data. Therefore, we gave individual identifiers to 32 slugs that were 7-days food-deprived and run in a group-feeding assay, recording whether each slug fed in a group or alone (GF, Fig 4A). In this first test, 13 of 24 animals fed socially. Six animals were removed from the analysis because they did not complete four of the subsequent tests.

**Fig. 4.**
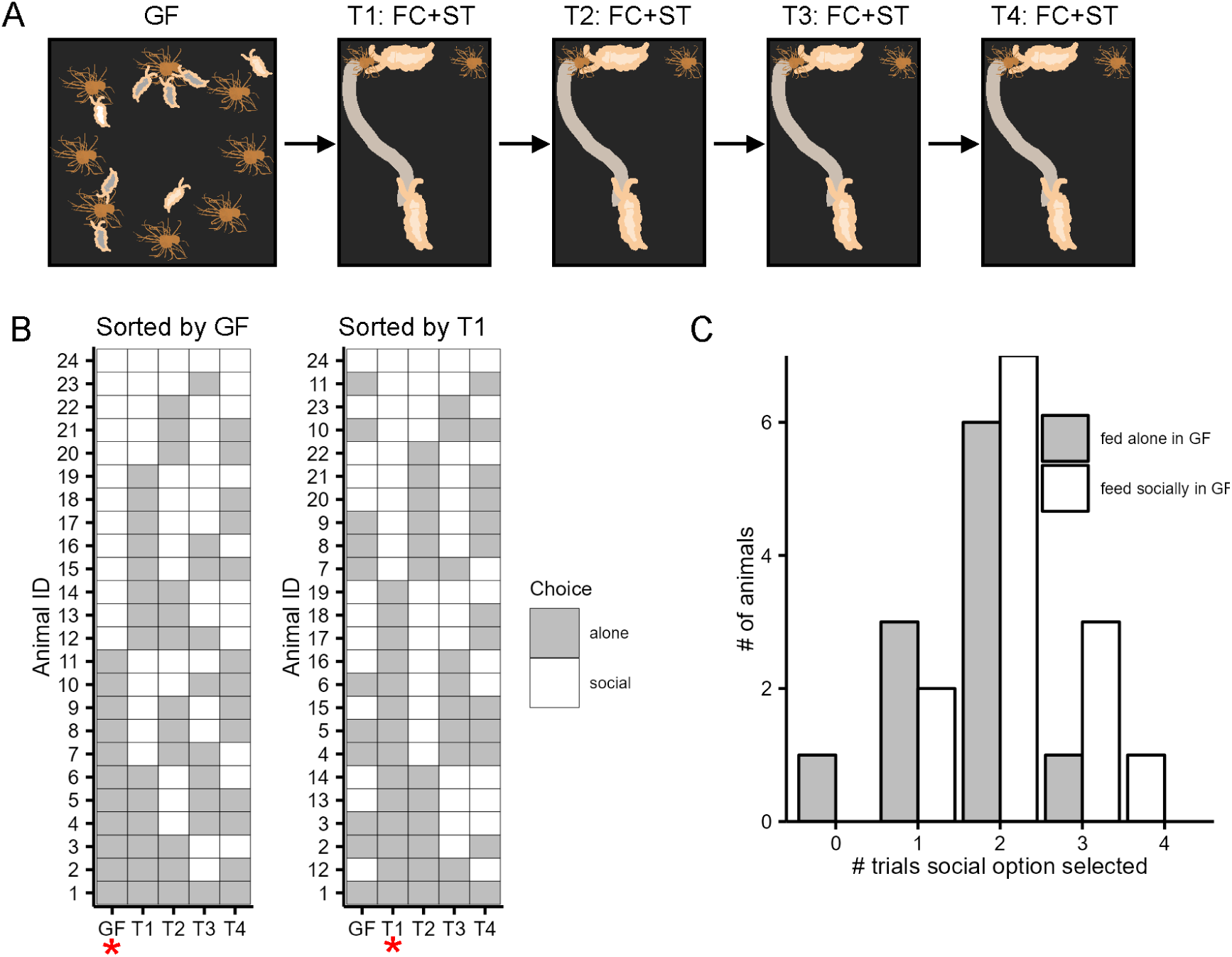
The choice to feed socially is not consistent within individuals. **A** Schematic showing the experimental design for this dataset. Animals were first tested in the group feeding assay (GF and then individually labeled and housed. Then they were tested 4 times (T1-T4) in a 2-alternative choice assay with a feeding conspecific and slime trail (FC+ST). **B** A plot showing the choices of each individual animal in the 5 different assays organized by their choice in the GF assay (left) and their choice in the T1 assay (right). **C** A histogram showing the number of animals that fed socially in the 2-alternative assays (T1-T4) 0 – 4 times. White represents animals that fed alone in the GF assay and grey represents animals that fed socially in the GF assay. The distribution is not bimodal and animals seem to randomly switch between feeding socially and alone.

After testing in the GF assay, the slugs were housed individually in clear plastic deli cups and underwent a 24-hour period of ad-libitum access to *E. diaphana* followed by 7-days of food-deprivation. They were then tested in the FC+ST 2-alternative choice assay and their choice was recorded (T1). Then, they were allowed to eat for 24-hours and then were food-deprived for another 7 days. This process was repeated such that each animal was tested four times (Fig. 4a). Their choices in the subsequent assays were used to create a score for each animal that represented the total number of times each individual chose the social option (anemone with a feeding conspecific and a slime trail).

Their first choice was compared to subsequent choices. In the first 2-alternative choice trial (T1) 10/24 of animals selected the social option and 8/13 of them had fed socially in the GF assay and 7/13 of them fed socially in the second 2-alternative choice trial (T2; Fig. 4b). The choice to feed socially in the GF assay was not predictive of how many times an animal would select the social option in the 2- alternative choice assays (Fisher’s Exact Test, p = 0.1548). If individual animals had consistent preferences to feed in groups, we would also expect a bimodal distribution in the number of trials they selected the social option, but the distribution was unimodal (Silverman (1981) critical bandwidth test, Critical bandwidth = 0.3612, p = 0.738; Fig. 4c). Their choices were not repeatable (R = 0, 95% confidence interval (CI) = 0., 0.136, p = 0.5).

## Discussion

Our study showed that *B. stephanieae* feed on their prey socially. These groups are larger than expected if each slug chose an anemone independently of each other. This social predation occurred with the same intensity in animals that were 7-days food-deprived and 3-days food-deprived, indicating that the level of satiety does not influence their propensity to feed in groups. We hypothesize that this social predation is a strategy to lessen the danger of injury from their radially symmetric prey by feeding from multiple sides. There are several other animals that show facultative social feeding behavior such as some species of otters that are mostly solitary except for bouts of cooperative hunting, which are thought to be in response to particularly fast prey or when other competitors for a prey item are high risk (Lodé et al., 2021). Similarly, sailfish often hunt individually but sometimes increase their prey capture rate by hunting schools of smaller fish together (Herbert-Read et al., 2016; Logan et al., 2023). The benefits of social predation vary across time and context for individuals, which maintains the use of multiple strategies (Lett et al., 2004; Macdonald, 1983). For example, African wild dog populations show differences in their level of cooperation and choreography during group hunting depending on their environment (Creel and Creel, 1995; Hubel et al., 2016). A key benefit of social predation is the capacity to subdue and kill larger, more dangerous prey (Brown and Richardson, 1988; MacNulty et al., 2014; Mukherjee and Heithaus, 2013). Since *E. diaphana* poses a threat to *B. stephanieae*, it is possible that their social predation is a strategy to avoid incapacitating injury that will lead to death.

The trade-offs of social foraging and social predation strategies have been studied using a combination of modeling and experimental approaches that determine different conditions under which social predation is likely to evolve. Hunting and foraging in groups can make it easier to find food because the load of searching and identifying a food source is split among more individuals (Snijders et al., 2021), however once located, the food must be shared across more individuals (Sutton et al., 2015). If animals experience predation risk, a benefit of social feeding is increased vigilance of the group with reduced energy spent on individual vigilance allowing more time to feed (Barta et al., 2004; Kelley et al., 2011). In fact, some animals increase aggregation behavior in the presence of odors from their predators, as in the freshwater amphipod *Gammarus pulex* (Kullmann et al., 2008). A down side is that groups can make it easier for predators to spot and capture individuals and this is a reason many hunters specifically target colonial prey (Balaban-Feld et al., 2019; Sutton et al., 2015).

Some individuals may inhabit temporary roles such as scrounging or producing where individuals either use the strategy of joining groups feeding at specific food sources or locate their own sources, respectively (Vickery, 2020). The choice of tactics is influenced by an individual’s early life experience (Katsnelson et al., 2008), perceived predation risk (Barta et al., 2004), hunger (Lendvai et al., 2004), and the availability and quality of food sources (Kurvers et al., 2012). Modeling studies support the idea that social predation can be maintained in populations where individuals may inhabit temporary roles such as scrounging or producing where individuals either use the strategy of joining groups feeding at specific food sources or locate their own sources, respectively (Vickery, 2020).

We sought to identify the cues that facilitated social predation in *B. stephanieae*. When tested in 2-alternative choice assays, *B. stephanieae* did not show a preference any of the cues related to the social option that we tested; they were not more attracted to slime trails, injured anemones or feeding conspecifics. This perplexing lack of preference could indicate some sort of density dependence where two slugs was not sufficient to drive the behavior, however in the group feeding assay many of the groups would be a pair of slugs and the first slug to join a group must have been attracted by cues from a single feeding slug.

In all of these assays, we also measured the size of the anemones and found that prey size did not impact likelihood of selecting the social option, which contrasts with findings in many vertebrate social predators (Hansen et al., 2023) and social spiders (Grinsted et al., 2020). This led to the hypothesis that individual slugs have different likelihoods of using social predation as a strategy. If the animals preferring social predation and animals that prefer to feed alone are random in the overall population of *B. stephaneiae*, then randomly sampling from the animals for the 2-alternative choice assay would show a null result. Many social predators have stable individual roles across hunting bouts such as the social spider *Australomisidia ergandros*; individuals specialize in a feeding tactic as a producer or a scrounger (Dumke et al., 2016). Similarly, individual dolphin specialize as divers and blockers when herding prey for capture (Gazda et al., 2005). This hypothesis also was not supported, indicating that individuals take on temporary roles as leaders and followers that drive fission-fusion social dynamics in the presence of prey. Each individual is not foraging randomly, however their roles seem to differ depending on context and specific foraging bout. This is similar to false cleaner fish, which have temporary roles when predating upon fish eggs (Sato et al., 2024) and the yellow saddle goat fish whose role is determined by spatial position in relation to the prey (Steinegger et al., 2018).

The 2-alternative choice assay may not be sufficient for identifying cues in social feeding because they capture only the initial attraction and choice. In the nematode, *Caenorhabditis elegans*, injury thus induces social feeding through activation of nociceptive neurons (de Bono et al., 2002). Since the 2-alternative choice assays were stopped at first contact between the slugs and their prey, it may not have allowed them to be injured and then re-evaluate their decision and ultimately select the other anemone. Individual *B. stephanieae* may need to interact with their prey for a longer time period and then be allowed to make a selection.

It is possible that the grouping behavior is a result of differences in the attractiveness of the prey, where in the group feeding assay slugs were attracted to the most attractive anemones and not to each other. This occurs in other systems, such as the way some individual humans are more attractive to mosquitos than others due to combinations of kairomones given off by a specific individual (Ellwanger et al., 2021; Giraldo et al., 2023). This differential attraction is seen at a species level where some humans have more frequent bites from multiple individual mosquitos. The differences in attraction seem to be associated with a variety of different host factors including diet, skin microbiome, levels of air-borne carboxylic acids and more. Similarly, some individual *E. diaphana* may have different sets of cues that make them more attractive. However, if these combinations of cues increased attraction in a way that was consistent across the species, then the assays that used a helper slug (Table 1) would still capture that effect and one would expect them to select the manipulated anemone at a rate higher than chance.

This study focused on adult behavior; however, unpublished observations of juvenile post-metamorphosis *B. stephanieae* also show social feeding behavior similar to adults (KO). In many animals, including this one, mixed sized individuals hinder individual growth. Juvenile *B. stephanieae* have higher mortality and lower growth rates when housed with adults (Monteiro et al., 2020). The relationship between developmental stages and the trade-offs associated with social foraging are often complicated and context dependent. For example, while juvenile ground squirrels forage in groups more, they do not reduce their vigilance like adults do in groups despite an increase in vigilance in solo foraging adults (Ortiz et al., 2019). In the algae-eating saccoglossan sea slug, *Placida dendritica*, feeding conspecifics stimulate others to feed and small slugs always benefit from foraging in groups, however large slugs only benefited with similarly sized conspecifics (Trowbridge, 1991). The specific pressures facing *Berghia* are likely to be complex and may be influenced by their developmental stage. Additionally, these experiments were conducted in a laboratory. The density of populations of this species in the wild and their feeding behaviors have not been studied. It is likely the social predation is dependent on the density of populations in the wild. Future work could investigate social predation in wild populations of *B. stephanieae*.

## Data Availability

The datasets generated and analyzed in the current study are available on Github (https://github.com/OtterLabGroup/nudibranch_social_predation_2024). Behavioral videos are available upon request.

## Author Contributions

All authors contributed to the study conception and design. Data collection and analysis were performed by Kate Otter and Saida Gamidova. The first draft of the manuscript was written by Kate Otter and all authors commented on previous versions of the manuscript. All authors read and approved the final manuscript.

## Acknowledgements

We thank the members of the Katz Lab for feedback on experimental design and for help with animal husbandry. We are grateful to the UMass Amherst Statistical Consulting (Yuijan Wu, Arthur Siller and Krista Gile) for their insight and assistance with selecting appropriate statistical models.

