## Supplementary Information for "Social predation by a nudibranch mollusc"

Electronic Supplementary Information

Behavioral Ecology and Sociobiology

Kate Otter, Saida Gamidova, Paul S. Katz

**Supplementary Table S1** ANOVA for latency to select an anemone during 2-choice alternative assays

| | Df | Sum squares | F-value | P-value |
|---|---|---|---|---|
| Choice | 1 | 0.165 | 0.165 | 0.293 |
| Assay Type | 3 | 3.930 | 8.751 | 1.89e-05 |
| Acclimation | 1 | 2.611 | 19.066 | 2.34e-05 |
| Choice: Assay Type | 3 | 0.274 | 0.617 | 0.605 |
| Choice: Acclimation | 1 | 0.803 | 5.864 | 0.0166 |
| Assay type: Acclimation | 1 | 0.100 | 0.734 | 0.3931 |
| Choice: Assay type: Acclimation | 1 | 0.227 | 0.346 | 0.1998 |

**Supplementary Table S2** Post-hoc comparisons of manipulations on latency to select an anemone during 2-alternative choice assays

| Manipulation | Comparison | Mean Difference | SE | df | p-value |
|---|---|---|---|---|---|
| BA | FC | -0.0696 | 0.0930 | 154 | 0.877 |
| | MA | 0.352 | 0.112 | 154 | 0.0105 |
| | ST | 0.113 | 0.091 | 154 | 0.604 |

| | | | | | |
|---|---|---|---|---|---|
| FC | MA | 0.422 | 0.100 | 154 | 0.0003 |
| | ST | 0.1823 | 0.077 | 154 | 0.085 |
| MA | ST | -0.239 | 0.099 | 154 | 0.0758 |

### Supplemental Fig. S1

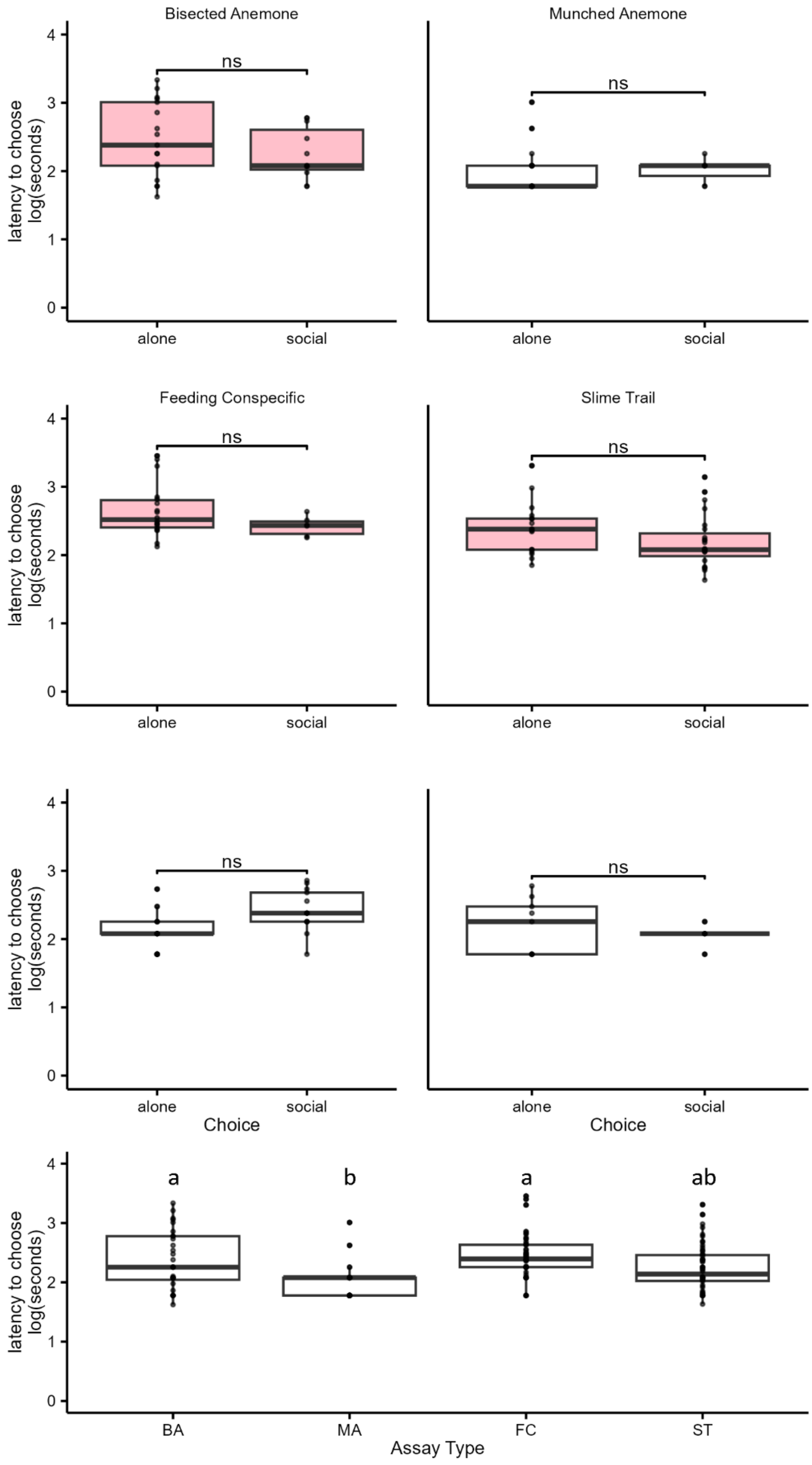

**Supplemental Figure S1** Animals that selected the social option in the 2-alternative choice assays did not choose faster than animals that did not select that option. The pink boxplots represent animals that were acclimated in ASW and the white boxplots represent animals that were acclimated in anemone-scented ASW. The data was log-transformed to normalize. The bottom boxplot shows the data with acclimation aggregated to show the results of pairwise post-hoc comparisons for the statistically significant predictor (assay type).

#### Supplemental Fig. S2

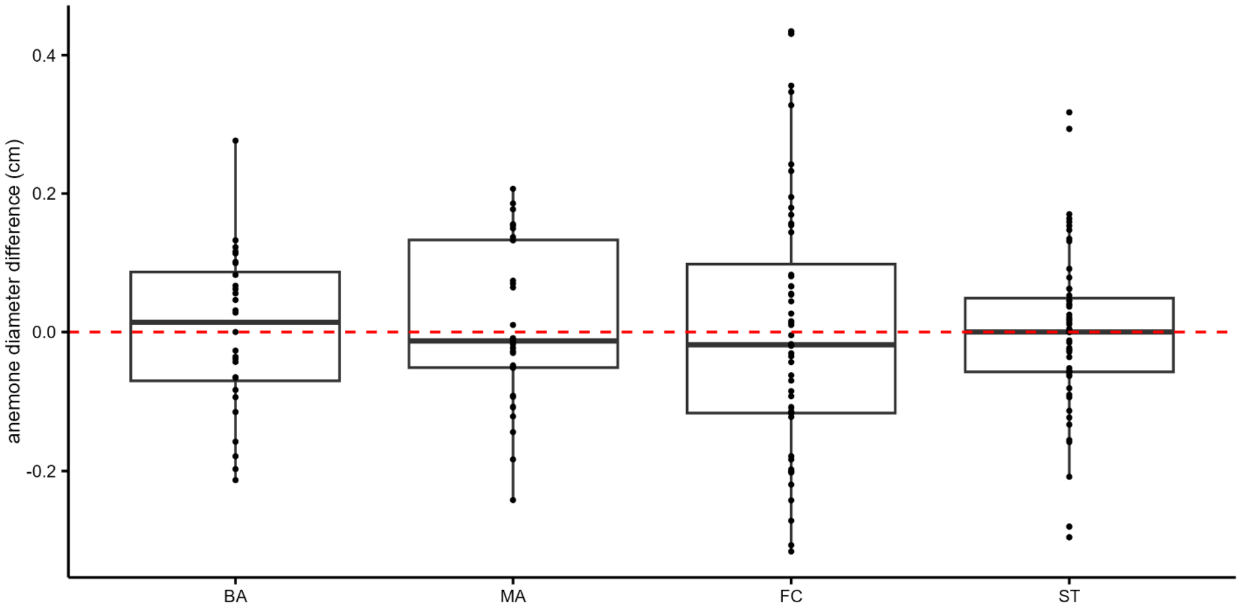


**Supplemental Figure S2** Animals did not consistently select larger or smaller anemones in the 2-alternative choice assays
